# Inhibition of Atrial Natriuretic Peptide Clearance Reduces Myocardial Fibrosis and Improves Cardiac Function in Diabetic Rats

**DOI:** 10.1101/2024.08.01.606125

**Authors:** Jules Joel Bakhos, Youakim Saliba, Joelle Hajal, Guy Achkouty, Hrag Oskaridjian, Chloé Azevedo, Albert Semaan, Nadine Suffee, Elise Balse, Stéphane N Hatem, Nassim Fares

## Abstract

**Background:** Natriuretic peptides (NP) exert pleotropic effects through the recruitment of cGMP-signaling pathways depending on their bioavailability which is regulated by clearance receptors and peptidases. Here, we tested the hypothesis that increasing myocardial bioavailability of NP has a beneficial effect on heart failure. We studied the effects of a mutated NP, MANP, resistant to neprilysin in a model of diabetic cardiomyopathy characterized by a marked myocardial fibrosis.

**Methods:** Natriuretic peptides as well as sacubritril were delivered via osmotic mini-pumps to high-fat/streptozotocin-induced type-2 diabetic (T2D) rats. Cardiac function was evaluated by echocardiography. Myocardial remodeling was studied by histological approaches, collagen phenotype and measurement of cGMP tissue concentration. Live-cell cGMP biosensing was conducted on cultured rat cardiac fibroblasts to investigate biological effects of NPs. cGMP signaling pathway was studied using various antibody arrays and biochemicals assays in cardiac tissue and cultured fibroblasts.

**Results:** MANP exhibits superior efficacy than ANP in reducing left ventricular dysfunction and to reduce myocardial fibrosis with less extracellular matrix deposition. *In vitro*, MANP and ANP similarly generated cGMP and activated PKG signaling pathway in cardiac fibroblasts, attenuating SMAD activation, collagen secretion and cell proliferation. Nevertheless, *in vivo*, MANP enhanced cardiac cGMP accumulation and was more potent than ANP in activating myocardial cGMP/PKG signaling and inhibiting the profibrotic SMAD pathway. Endopeptidase inhibition using sacubitril also led to cardiac cGMP accumulation and reduced myocardial fibrosis

**Conclusions:** Myocardial bioavailability of ANP is a major determinant of peptide efficacy in reducing cardiac fibrosis and improving pump function during diabetic cardiomyopathy.

**Clinical Perspective:** *What Is New?:*

- Mutated atrial natriuretic peptide (MANP) resistant to neprilysin degradation outperforms wild-type ANP in reducing myocardial fibrosis and improving cardiac function in type-2 diabetes (T2D)
- While the antifibrotic effect of the two ANP isoforms involves similarly cGMP-dependent PKG signaling and inhibition of fibroblast activation, MANP enhanced cGMP myocardial concentration more importantly than ANP.
- Sacubitril that inhibits ANP degradation also reduces cardiac fibrosis through myocardial accumulation of cGMP and activation of cGMP-dependent PKG signaling pathway.
- Cardiac bioavailability of natriuretic peptides is a major determinant of their effects on myocardial fibrosis and cardiac function.

*What Are the Clinical Implications?:*

- Myocardial bioavailability of natriuretic peptides is crucial for mitigating cardiac fibrosis and improving cardiac function in diabetic cardiomyopathy and heart failure in general.
- MANP holds the potential as a new treatment modality in the management of heart failure.

## Introduction

Natriuretic peptides (NP) initially recognized to regulate sodium and volume homeostasis and, hence, protect against high blood pressure, also show a range of beneficial effects on the heart, vessels, and kidney. They can also improve glucose tolerance and regulation of adipose tissue accumulation and biological properties ^1,2^. The pleiotropic effects of NP are primarily mediated by cGMP second messengers that are recruited following NP binding on type A- or type B-plasma membrane NP receptors which are widely expressed. Furthermore, biological effects of natriuretic peptides depend on their half-life, three minutes for atrial natriuretic peptide (ANP) and twenty minutes for brain natriuretic peptide (BNP), which is regulated by clearance receptor (NPR-C) and a variety of peptidases, including neprilysin ^3^.

Various pharmacological strategies have been developed to modulate the NP system, notably in the context of heart failure, characterized by altered cGMP pathway ^4,5^. For instance, human recombinant natriuretic peptides such as carperitide (for ANP) and nesiritide (for BNP), have been investigated for heart failure treatment, with, however, controversial clinical outcomes ^6^. Besides, another strategy has been pioneered by decreasing ANP elimination through neutral endopeptidase neprilysin inhibition. This leads to enhanced bioavailability of ANP that otherwise required continuous infusion of the peptide at a low dose ^3^. Sacubitril-Valsartan is the first-in-class agent combining neprilysin inhibitor and angiotensin-receptor blocker that has been shown to improve cardiovascular outcomes, myocardial performance and reduced hospitalization and mortality of heart failure patients ^7–11^. In addition to NP, neprilysin hydrolyzes several hormones such as bradykinin, angiotensin I and II, and endothelin that could contribute to the systemic and local effects of enzyme inhibition on cardiovascular and kidney axes ^12^. Undoubtedly, the precise pharmacological properties and mechanisms of action of NP as well as related pharmacological molecules remain to be determined.

One current hypothesis stipulates that depending on their tissue bioavailability, NP can have marked local tissular effects contributing to the reduction of myocardial hypertrophic and fibrotic remodeling ^1,2^. This hypothesis was tested here using a mutated isoform of ANP - MANP-resistant to neprilysin degradation ^13,14^ with a superior bioavailability and efficacy as to ANP ^15,16^. Not only is MANP resistant to neprilysin degradation but to insulin degrading enzyme (IDE) also with an IC50 that is 30-fold higher than ANP ^13,14^. Indeed, MANP is actively under clinical investigation for resistant and essential hypertension treatment ^17–20^. Herein, we used the high-fat/streptozotocin-induced type-2 diabetes (T2D) model that reproduces the diabetic cardiomyopathy notably characterized by diastolic dysfunction and marked myocardial fibrosis ^21–26^. We provide novel evidence that modulating peptide bioavailability determines the effects of ANP on cardiac fibrosis.

## Methods

### Ethical statement

The present research was approved by Saint Joseph University of Beirut’s Ethical Committee (Project #FM348, mixed project USJ/CNRS-L). The protocols used were developed in accordance with the American Physiological Society Council’s Guiding Principles in the Care and Use of Animals, the US National Institutes of Health’s Guide for the Care and Use of Laboratory Animals (NIH Publication no. 85-23, revised 1985), and the European Parliament Directive 2010/63 EU. The animals were housed in a controlled environment with a constant temperature of 25°C and humidity level of 50 ± 5%, and were exposed to a 12:12h light/dark cycle. They were given regular rodent food and unfettered access to tap water. They were given at least one week to adjust to these conditions before the inquiry began.

### Diabetes animal model and treatments

The study was carried out on male Wistar rats aged 12 months. The conventional high-fat diet combined with streptozotocin (STZ) (Sigma-Aldrich, St Louis, MO, USA) injections was used to induce T2D. Mice were fed a high-fat diet (60% kcal from lard) for four weeks before receiving three intraperitoneal streptozotocin injections (40 mg.kg^-1^) on three consecutive days. STZ was diluted with a citrate buffer at pH 4.5. Sham animals received citrate injections. To guarantee proper diabetes induction, blood glucose levels were measured 72 hours following the third STZ injection. After a 12-hour fast, conscious rats were placed in rodent plexiglass restrainers (IITC Life Science Inc., CA, USA), and blood glucose levels were measured from the tail using a glucometer (Accu-Chek, Roche Diabetes Care, IN, USA). The tails were first cleaned with 70% alcohol, and measurements were taken with the second drop of blood. Diabetic animals had blood glucose levels higher than 200 mg.dl^-1^. Under ketamine/xylazine (50 and 10 mg.kg^-1^; Panpharma, Luitré, France and Interchemie, Waalre, Holland) anesthesia, osmotic mini-pumps (model 2004; Alzet, Durect, CA, USA) were implanted subcutaneously on the back posterior to the scapulae. The pumps delivered 0.25 µl.hr^-1^ and a dosage of 2 pmol.Kg^-1^.min^-1^ of ANP or MANP (AnaSpec, Fremont, CA, USA) diluted in sterile 0.9% NaCl. Prior research has shown that these concentrations have little effect on systemic hemodynamics ^15^. Sacubitril sodium (MedKoo Biosciences Inc., Durham, NC, USA) was administered in drinking water at a dose of 18 mg.kg^-1^.day^-1^, corresponding to a human equivalent dose (HED) of 2.85 mg.kg^-1^.day^-1^ (≈ 97mg sacubitril from the Entresto® twice daily recommended maintenance dose for an adult weighing 70 kg). The animals were randomly assigned to two independent sets of experiments, the first set consisting of four groups (n=6 each): Sham, T2D, T2D ANP, T2D MANP, and the second set consisting of five groups: Sham (n=7), T2D (n=6), T2D ANP (n=5), T2D SACU (sacubitril) (n=7), T2D ANP SACU (sacubitril) (n=6).

### Echocardiography

The high-resolution color Doppler ultrasound system SonoScape S2V (SonoScape Co., Shenzhen, China) with a 9 MHz C611 transducer intended for mice and rats was used to perform transthoracic echocardiography. An EZ-SA800 Anesthesia Single Animal System (E-Z Systems, Pennsylvania, USA) was used to anesthetize the rats prior to sacrifice. The dose of isoflurane (Baxter, Deerfield, IL, USA) used was 5% for induction and 3% for maintenance, with a flow rate of 1 L/min. The parasternal long-axis 2D view of the left ventricle was captured in M-mode at the level of the papillary muscles to assess the thickness of the ventricular walls and internal diameters, enabling calculations of fractional shortening (FS) and ejection fraction (EF) using the Teichholz method. This procedure was carried out by two independent operators who were unaware of the experimental conditions. All measurements were taken between 9 a.m. and noon, with duplicate readings acquired for each condition.

### Rat ventricular cardiac fibroblasts isolation and culture

Wistar rats were anesthetized with ketamine/xylazine, the depth of anesthesia was checked by the pedal withdrawal reflex, then hearts were excised and ventricles placed in ice-cold modified Tyrode solution with the following composition (in mM): 117 NaCl, 5.7 KCl, 1.7 MgCl_2_, 4.4 NaHCO_3_, 1.5 KH_2_PO_4_, 10 HEPES, 10 creatine monohydrate, 20 taurine, 11.7 D-glucose, 1% bovine serum albumin, adjusted to pH 7.1 with NaOH. Ventricles were minced and subjected to four rounds of enzymatic digestion using modified Tyrode solutions: collagenase type V (165.1 U/ml) and protease type XXIV (4.62 U/ml) from Sigma-Aldrich, St. Louis, MO, USA, for 20 minutes at the beginning, and collagenase type V (157 U/ml) for three more 20-minute rounds. Supernatants from each bath were pooled and enzymatic reaction stopped by adding cold fetal bovine serum. After centrifugation at 500 rpm for 10 minutes, the cardiomyocytes were eliminated, and the non-myocyte cells were separated by centrifugation at 2000 rpm for 10 minutes. The cells were then suspended in Dulbecco’s Modified Eagle’s Medium (DMEM), supplemented with 1% penicillin/streptomycin and 10% heat-inactivated fetal bovine serum from Sigma-Aldrich, St. Louis, MO, USA. After incubating for two hours, non-adherent cells were discarded, and the remaining adherent fibroblasts were rinsed with fresh culture medium. Cells were counted and equally dispatched to maintain consistency across conditions. After 24 hours, cells had the fibroblast-like fusiform and spindle morphology. Cells were studied after 5 days of culture. A subset of cells was treated for 24 hours with high glucose to mimic hyperglycemia with or without ANP and MANP (100 pM).

### cGMP live-cell imaging

Live cultured cardiac fibroblasts were used to monitor cGMP synthesis. This was done by utilizing a Genetically Encoded Nucleotide Indicator (GENIe) that carried mNeonGreen, a highly luminous monomeric green fluorescent protein. GENIe (Montana Molecular, MT, USA) is a sensor that exhibits a strong initial fluorescence which decreases as the intracellular cGMP levels increase. Experiments were conducted following the manufacturer’s protocol. After cell synchronization on day 3 of culture, the cells were exposed to a BacMam modified baculovirus along with sodium butyrate, an HDAC inhibitor that is essential to sustaining viral expression. After 24 hours of infection, the cell culture medium was replaced and the cells were left for an additional 24 hours. The cells were placed in the modified Tyrode solution and left to incubate for 15 minutes prior to starting the recordings. Cells were acutely perfused with the different peptides using a ValveLink8.2 Perfusion Controller (Automate Scientific, CA, USA). Images were acquired and analyzed using a digital fluorescent imaging system (InCyt Im2; Intracellular Imaging Inc., Cincinnati, OH, USA).

### Cardiac and plasma cGMP measurement

Blood was collected from each animal group into EDTA tubes and centrifuged at 4500 rpm for 10 minutes to separate plasma. cGMP plasma concentrations were measured using ELISA kits according to the manufacturer’s instructions (Cyclic GMP Complete ELISA Kit; ab133052; Abcam, Cambridge, United Kingdom). To measure ventricular cGMP concentrations, cardiac tissues were first homogenized in 0.1 M HCl before performing the test. Acetylation of the samples was carried out to improve the sensitivity of the cGMP assay.

### Antibody array and Western blot

Ventricular tissue and cultured cardiac fibroblasts were homogenized and lysed in RIPA assay lysis buffer including protease and phosphatase inhibitors. Protein concentrations were measured using the Bradford protein assay (Bio-Rad, Marnes-la-Coquette, France), and samples were denatured in Laemmli loading buffer (Bio-Rad, Marnes-la-Coquette, France) containing 10% β-mercaptoethanol (Sigma-Aldrich, St. Louis, MO, USA) at 37°C for 20 minutes. Proteins were separated by SDS 12% PAGE and transferred to polyvinylidene fluoride membranes (Bio-Rad, Marnes-la-Coquette, France) blocked with 5% bovine serum albumin. The membranes were incubated overnight at 4°C with the corresponding primary antibodies: anti-PKG-I (protein kinase G) (ab90502; 1/1000), anti-VASP (vasodilator-stimulated phosphoprotein) (ab209093; 1/1000), anti-phospho-VASP Ser239 (ab194747; 1/500), anti-SMAD2 (Mothers against decapentaplegic homolog 2) (ab63576; 1/500), anti-phospho-SMAD2 S467 (ab53100; 1/500), anti-SMAD3 (Mothers against decapentaplegic homolog 3) (ab28379; 1/500), and anti-phospho-SMAD3 S423+S425 (ab52903; 1/500). Blots were re-probed for glyceraldehyde-3-phosphate dehydrogenase (GAPDH) (#5174; 1/2500) from Cell Signaling Technology, Danvers, MA, USA. Goat anti-rabbit (1/3000, Bio-Rad Laboratories, Marnes-la-Coquette, France) was used as the secondary antibody. An imaging system fitted with a CCD camera was used to detect enhanced chemiluminescence signals (Omega Lum G, Aplegen, Gel Company, SF, USA). Image Studio Lite Ver 5.2 (LI-COR Biosciences, Lincoln, NE, USA) was used to do the quantifications. For each condition, three western blots were used.

To generate antibody arrays, 5 µg of proteins were immobilized onto PVDF membranes using dot blot apparatus (Cleaver Scientific, Rugby, UK) before blocking and incubating with the primary antibody. The same GAPDH and secondary antibodies were utilized as in the western blot studies with the additional anti-Col1 (ab34710; 1/1000) and anti-Col3 (ab7778; 1/1000) (Abcam).

### Histology and immunofluorescence

Formalin-fixed cardiac tissues were embedded in paraffin, cut into 4 μm slices, and stained with Masson’s Trichrome or Sirius red (Sigma-Aldrich, St. Louis, Missouri, USA). Histological examinations were conducted by two independent pathologists, and fibrosis was assessed using a scoring procedure, an Aperio Slide Scanner and ImageScope software v12.4.6.5003 (Leica Biosystems, Wetzlar, Germany), and a VanGuard High-Definition Digital Camera (VEE GEE Scientific, Illinois, USA). WGA staining was carried out in accordance with the manufacturer’s instructions (Thermo Fischer Scientific, MA, USA). PDGFRα labeling was followed by antigen retrieval with HCl. Triton X-100 was used to permeabilize the membrane, and 10% goat serum and 1% bovine serum albumin were used for blocking. PDGFRα antibody (ab124392; 1/100; Abcam, Cambridge, USA) was employed overnight at 4°C to detect fibroblasts. The secondary antibody was goat anti-rabbit IgG H&L Alexa Fluor 594 (ab150084; Abcam), which was pre-adsorbed in a column matrix containing immobilized mouse serum proteins to reduce background by minimizing cross-reactivity with endogenous proteins and immunoglobulins. Sections were mounted with Fluoroshield Mounting Medium containing DAPI (Abcam, Cambridge, UK), and images were taken with an Axioskop 2 immunofluorescence microscope (Carl Zeiss Microscopy GmbH, Jena, Germany) outfitted with a CoolCube 1 CCD camera (MetaSystems, Newton, Massachusetts, USA). Images were analyzed using ImageJ.

### Soluble collagen and proliferation assays

MTT assay was used to analyze cell proliferation. The cell culture medium was withdrawn, and 0.5 mg.ml-1 MTT Tyrode solution was added to the cells. After an hour of incubation at 37°C, the MTT solution was discarded and the cells were rinsed with Tyrode. MTT formazan purple crystals were dissolved in 100% DMSO, and absorbance was measured at 550 nm.

The Sircol collagen test (Biocolor, County Antrim, UK) was used to measure soluble collagen. In brief, ventricular tissues were homogenized in an acid-pepsin solution, whereas cultured cell supernatants were used directly. Each sample and standard were then treated with the Sircol dye reagent. Collagen-dye pellets were centrifuged, rinsed with acetic acid, and then dissolved in an alkali reagent. Absorbance was measured at 550 nm.

### Statistical analysis

All quantitative data are reported as mean ± SEM with individual data points. Normal distribution and equal variance of the values were verified using the Shapiro-Wilk and Bartlett’s tests, respectively. Statistical analyses included Repeated-measures two-Way ANOVA followed by Sidak’s test, One-way ANOVA followed by Sidak’s or Holm-Sidak’s test, Kruskal-Wallis with Dunn’s test, and Brown-Forsythe and Welch ANOVA followed by Dunnett’s T3. Statistical significance was denoted by *P<0.05, **P<0.01, ***P<0.001, and ****P<0.0001.

## Results

### MANP exhibits superior efficacy than ANP in reducing left ventricular dysfunction and fibrosis associated with T2D

In male rats subjected to streptozotocin injections coupled to high-fat diet for six weeks, we compared the effects of low doses of ANP and MANP delivered subcutaneously by osmotic pumps on cardiac remodeling and dysfunction (Figure 1A). Three days after diabetes induction, animals showed high fasting plasma glucose (>200mg.dl^-1^) as compared to sham (<100mg.dl^-1^). A further increase of fasting plasma glucose was noted at week 6 (≈500mg.dl^-1^). Similar plasma glucose patterns were observed in the other groups albeit fasting plasma glucose remained unchanged at week 6 under ANP and MANP (Figure 1B). Cardiac mass was unaltered when normalized to body weight in all groups (Figure 1C). Echocardiographic analyses showed a trend towards LV dilation (Figure 1D-1F) in T2D animals. As expected, left ventricular function was declined in T2D (EF≈75%, FS≈37% in T2D *vs* EF≈95%, FS≈63% in sham). Left ventricular dysfunction was moderately improved under ANP (EF≈82%, FS≈44%), and almost restored with MANP treatment (EF≈92%, FS≈56%) (Figure 1G, 1H). Microscopically, T2D rats developed diffuse myocardial fibrosis depicted by Masson’s trichrome with an expansion of extracellular matrix labelled by wheat germ agglutinin (WGA) and spread of activated PDGFRα^+^ cardiac fibroblasts (Figure 1I-1K). MANP treatment showed superior efficacy than ANP in protecting rat hearts with ≈80% and 75% decrease in Masson’s trichrome and WGA staining respectively as compared to ≈50% decrease with ANP (Figure 1I, 1J). In contrast, similar fibroblast labelling extent was noted under both peptides (Figure 1K). At the molecular level, MANP inhibited SMAD3 activation as indicated by ≈55% lower phosphorylation levels compared to T2D animals (Figure 1L). These results indicate that MANP more efficiently than ANP reduces cardiac dysfunction and myocardial fibrosis in T2D rats. Subsequently, cardiac collagen characterization was done by antibody arrays for the major two collagen isoforms, collagen I and III as well as Sircol assay for newly synthesized soluble collagen. Cardiac collagen III but not collagen I increased in T2D (by ≈55%) coupled to a decrease in soluble collagen (by ≈30%), whereas ANP-treated groups showed opposite cardiac collagen profiles (decrease by ≈30% in collagen III and increase by ≈25% in soluble collagen). Of note, MANP led to significant decreases in both forms of collagen (by ≈50% in collagen III and by ≈20% in soluble collagen) (Figure 1M-1O).

**Figure 1.**
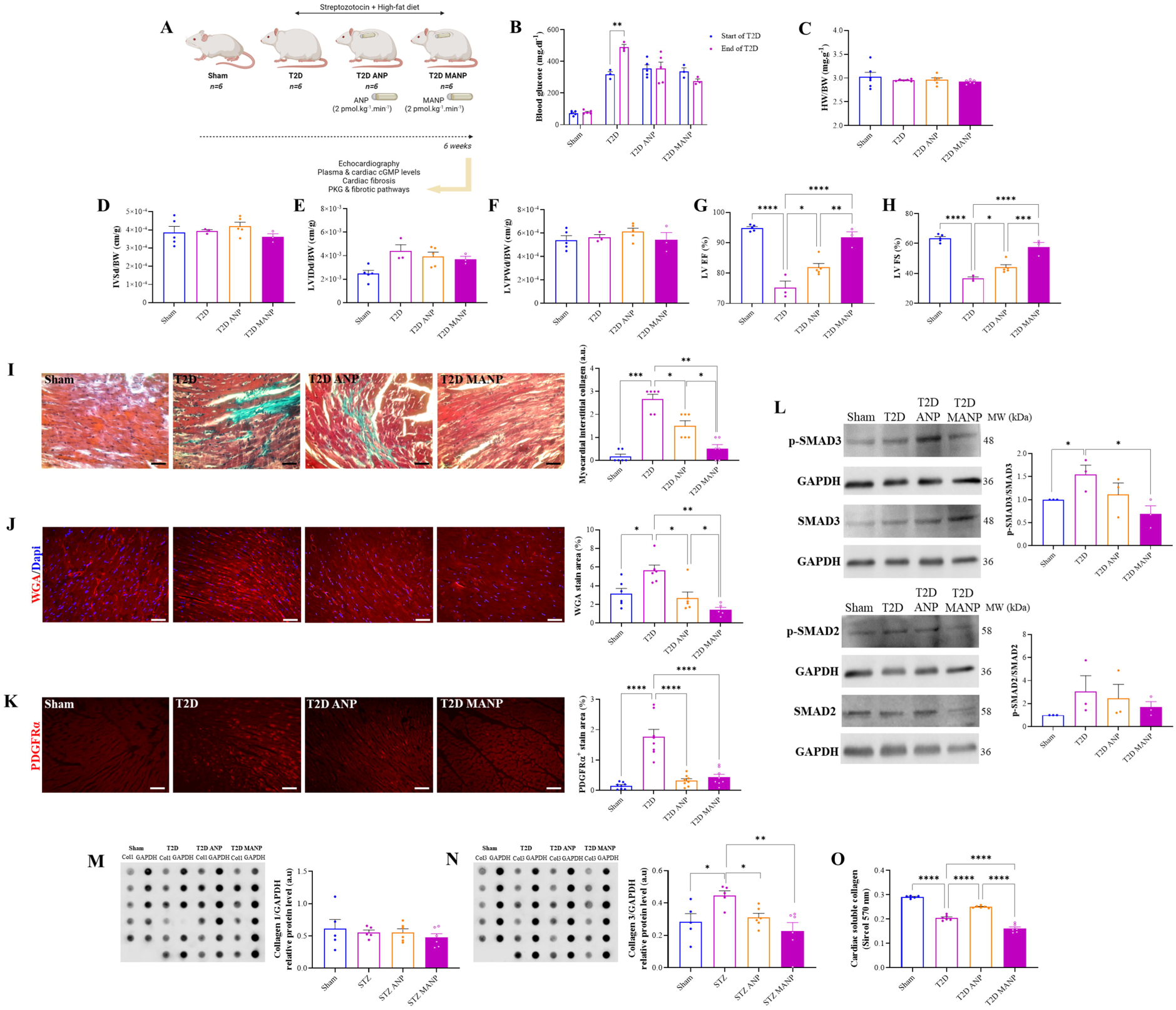
Left ventricular remodeling in T2D is mitigated by MANP more effectively than ANP. **A,** Experimental design for subjecting male rats to streptozotocin injections coupled to a high-fat diet as a model for T2D. ANP/MANP were delivered by subcutaneous osmotic pumps at a non-diuretic dose of 2 pmol.Kg^-1^.min^-1^. Cardiac function was assessed by echocardiography at six weeks of the protocol. Rats were then euthanized, and hearts and plasma were harvested for further histological and molecular analyses. Echocardiographic parameters were normalized to body weight to account for body size variation among the rats and interpret measurements independently of the individual animal’s size. **B,** Representative blood glucose in Sham, T2D, T2D ANP, and T2D MANP. **C,** Mean heart weight over body weight ratio (HW/BW), **D,** mean interventricular septum thickness at end-diastole, **E,** mean left ventricular internal dimension at end-diastole, **F,** mean left ventricular posterior wall thickness at end-diastole, **G,** mean left ventricular ejection fraction, and **H,** mean left ventricular fractional shortening in all groups. **I-K,** Representative heart sections for all groups showing Masson’s trichrome staining, wheat germ agglutinin (WGA) and PDGFRα immunofluorescence (594 nm; bar = 50µm) with respective quantifications of analyzed section areas. **L,** Representative western blots and densitometric analyses of phosphorylated and total SMAD2/3 (each with GAPDH as housekeeper protein) for all groups. **M, N,** Antibody arrays for cardiac collagen I and III using GAPDH as internal control with respective quantifications for all groups. **O,** Cardiac soluble collagen for all groups using Sircol assay. T2D: type 2 diabetes; ANP: atrial natriuretic peptide; MANP: M-atrial natriuretic peptide. All quantitative data are reported as mean and individual data points ± SEM. Normal distribution of the values was checked by the Shapiro-Wilk test and equal variance was checked by the Bartlett’s test. Repeated-measures two-Way ANOVA followed by Sidak’s test in (**B**), One-way ANOVA followed by Sidak’s test in (**C-G, K-M, O**) or Holm-Sidak’s test in (**H, N**), Kruskal-Wallis followed by Dunn’s test in (**I**), and Brown-Forsythe and Welch ANOVA followed by Dunnett’s T3 in (**J**) were performed. \**P*<0.05, \*\**P*<0.01, \*\*\**P*<0.001, and \*\*\*\**P*<0.0001.

### ANP and MANP similarly regulate cyclic GMP signaling pathway in cardiac fibroblasts

As the antifibrotic effect of ANP has been attributed mainly to its action on cardiac fibroblasts ^27–29^, we examined whether difference in anti-fibrotic effect between ANP and MANP could be related to distinct effects on fibroblasts. The mNeon Green GENIe system, delivered through BACMAM, was employed for cyclic GMP biosensing in primary cultured cardiac fibroblasts. This downward-sensing system exhibits bright initial green fluorescence, which diminishes in response to increased intracellular cGMP levels. Upon 100 pM MANP or ANP acute addition, the biosensor’s fluorescence gradually decreased with comparable F/F_0_ for both peptides (Figure 2A, 2B). In addition, MANP and ANP similarly decreased rat cardiac fibroblast proliferation (by ≈20%) and collagen secretion (by ≈30%) as demonstrated by MTT and sircol assays (Figure 2C, 2D). Fibroblasts were subsequently subjected to a 24-hour treatment with elevated glucose, replicating conditions of diabetic hyperglycemia, w/wo MANP or ANP. Pro-fibrotic SMAD2 was activated by high glucose but showed decreased phosphorylation levels with both MANP and ANP (by ≈55%) whereas SMAD3 remained stable (Figure 2E). High glucose inhibited cGMP-dependent pathway PKG/VASP as depicted by lower VASP phosphorylation levels at serine 239, whereas MANP/ANP activated this pathway in a comparable manner (≈2.3 fold) (Figure 2F). Therefore, ANP and MANP show the same biological properties on cardiac fibroblasts.

**Figure 2.**
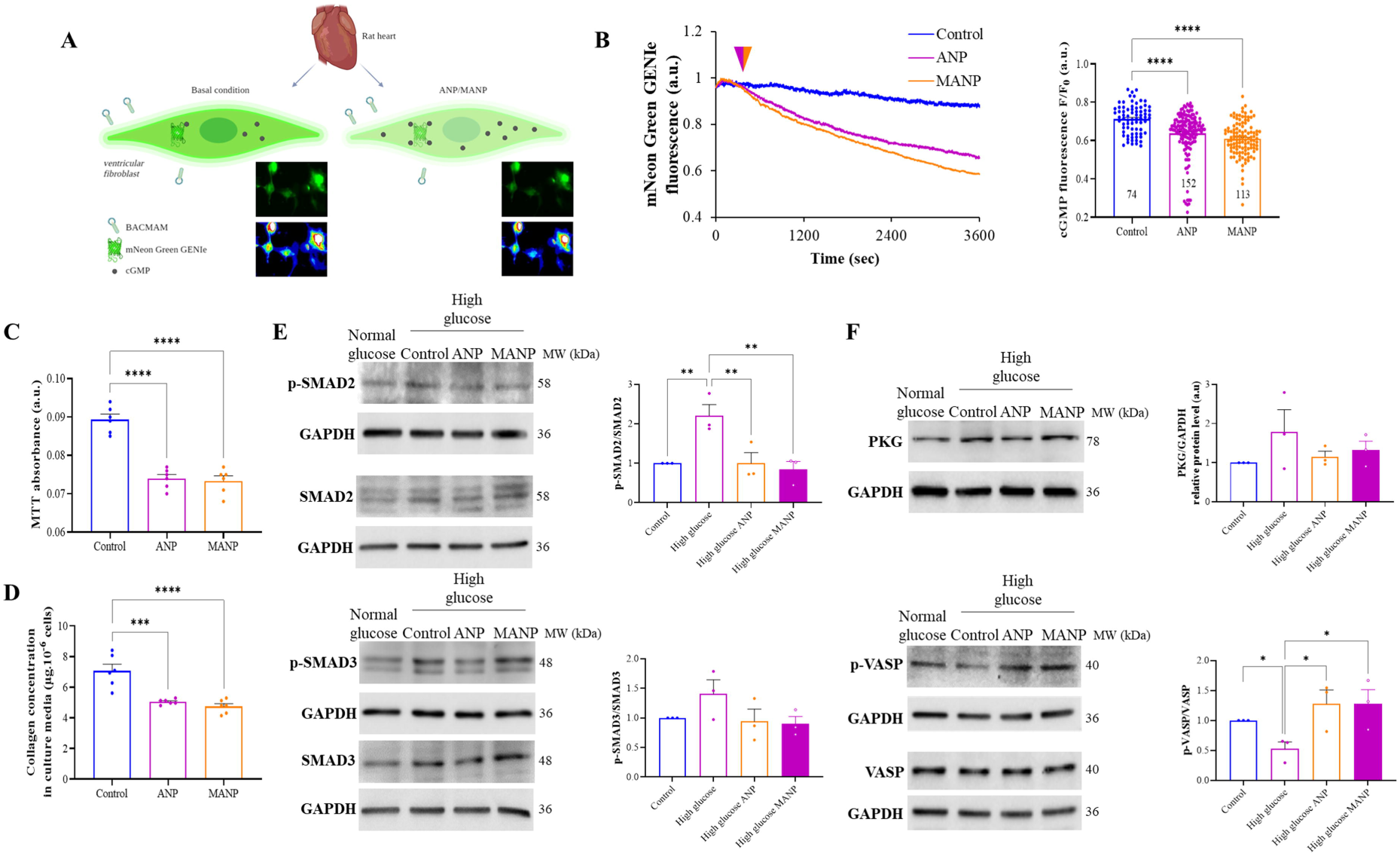
MANP-driven PKG activation and SMADs regulation in cardiac fibroblasts through cyclic GMP. **A,** Illustration of the mNeon Green GENIe system introduced through BACMAM for monitoring intracellular cGMP in live primary cardiac fibroblasts. The emitted green light from the probe exhibits an inverse relationship with cGMP concentration. MANP or ANP was acutely introduced to the cells during imaging at a concentration of 100 pM. Cells were isolated from normal rat hearts. **B,** Representative mNeon Green GENIe probe fluorescence curves after the acute addition of MANP or ANP and F/F_0_ fluorescence quantifications. **C, D,** Proliferation and collagen secretion (µg.10^-3^ cells) of cultured rat cardiac fibroblasts treated for 24 hours with 100 pM MANP or ANP, estimated by MTT and Sircol assays; absorbance at 570 nm. **E, F,** Cultured rat cardiac fibroblasts were treated with high glucose for 24 hours mimicking diabetic hyperglycemia w/wo MANP or ANP. Representative western blots and densitometric analyses of phosphorylated and total SMAD2/3 and vasodilator-stimulated phosphoprotein (VASP), and protein kinase G (PKG) (each with GAPDH as housekeeper protein) are shown for all groups. ANP: atrial natriuretic peptide; MANP: M-atrial natriuretic peptide. All quantitative data are reported as mean and individual data points ± SEM. Normal distribution of the values was checked by the Shapiro-Wilk test and equal variance was checked by the Bartlett’s test. One-way ANOVA followed by Tukey’s test in (**B-D**) or Holm-Sidak’s test in (**E, F lower panel**), and Brown-Forsythe and Welch ANOVA followed by Dunnett’s T3 in (**F upper panel**) were performed. \**P*<0.05, \*\**P*<0.01, \*\*\**P*<0.01, and \*\*\*\**P*<0.0001.

### MANP is more potent than ANP to activate cGMP/PKG signaling in the heart of T2D rats

We next examined whether differences in antifibrotic effect between wild-type and mutant ANP could be related to differences in peptide bioactivity, in line with the reduced clearance of the mutant peptide. Myocardial concentration of cGMP was measured to evaluate the activation of membrane particulate guanylate cyclase which is triggered by natriuretic peptides. Cardiac but not plasma cGMP levels presented a tendency to decrease (≈50%, *P*=0.13) in T2D which was attenuated in rats treated by ANP. In contrast, MANP treatment increased cardiac cGMP levels compared to all groups of rats (≈5 fold vs T2D) as well as plasma cGMP albeit not as important as within the heart (≈2.9 fold vs T2D) (Figure 3A, 3B). Furthermore, T2D inhibited cardiac cGMP-dependent pathway PKG/VASP as evidenced by lower VASP phosphorylation levels (by ≈60%); both MANP and ANP treatments restored this pathway (Figure 3C). These results are consistent with a more accumulation of MANP than ANP in the myocardium of T2D rats.

**Figure 3.**
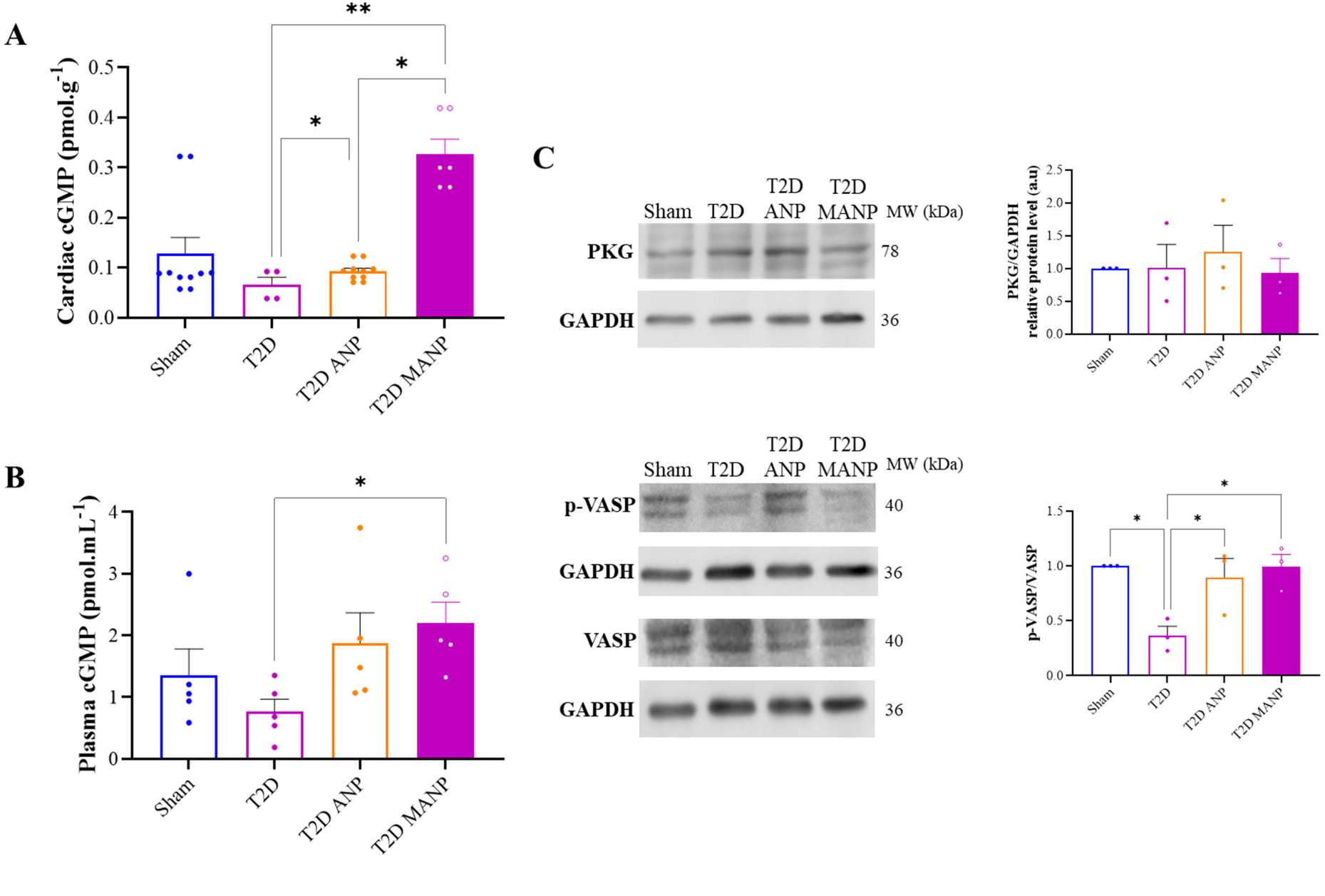
High cardiac sustainability of MANP restores cardiac cGMP/PKG signaling in T2D. **A, B** Plasma and cardiac cGMP concentrations measured by ELISA in the different rat groups. **C,** Western blots and quantifications of cardiac PKG, total and phosphorylated VASP in the different rat groups, with GAPDH as an internal control (n=3 for each condition). T2D: type 2 diabetes; ANP: atrial natriuretic peptide; MANP: M-atrial natriuretic peptide. All quantitative data are reported as mean and individual data points ± SEM. Normal distribution of the values was checked by the Shapiro-Wilk test and equal variance was checked by the Bartlett’s test. Kruskal-Wallis followed by Dunn’s test in (**A**), One-way ANOVA followed by Sidak’s test in (**B, C lower panel**), and Brown-Forsythe and Welch ANOVA followed by Dunnett’s T3 in (**C upper panel**) were performed. \**P*<0.05, and \*\**P*<0.01.

### Endopeptidase inhibition using sacubitril reduces myocardial fibrosis in T2D rats

To further examine the impact of natriuretic peptide accumulation level in the myocardium on its anti-fibrotic effects, we inhibited the endopeptidase neprilysin using sacubitril (Figure 4A). First, T2D rats treated with sacubitril showed the same metabolic profiles as their counterparts (Figure 4B). Following diabetes induction, cardiac mass was unaltered when normalized to body weight in all groups (Figure 4C) and there was no significant cardiac dilation in all groups of rats (Figure 4D-4F). Left ventricular function was reduced in T2D (EF≈78%, FS≈40%), and partially restored in ANP (EF≈85%, FS≈50%) and sacubitril (EF=87%, FS≈50%) conditions, without an additional effect of the association of ANP and sacubitril (EF≈83%, FS≈44%) (Figure 4G, 4H). Comprehensive analysis of myocardial interstitial fibrosis was conducted using Sirius red staining and showed elevated density of thickened collagen fibers in T2D (≈2 fold), that were significantly attenuated by both ANP and sacubitril treatments (by ≈65%, ≈50%, and ≈50% respectively, Figure 4I). Cardiac collagen was characterized using antibody arrays for collagen I and III, along with Sircol assay for soluble collagen synthesis. In T2D, collagen III increased by ≈55%, while soluble collagen decreased by ≈40%. ANP/sacubitril-treated groups exhibited opposite trends (≈30% decrease in collagen III, ≈30% increase in soluble collagen, Figure 4J-4L). Finally, the increased SMAD3 but not SMAD2 activation in diabetic hearts was diminished by ANP/sacubitril treatments with ≈65% lower phosphorylation levels as compared to T2D (Figure 4M, 4N).

**Figure 4.**
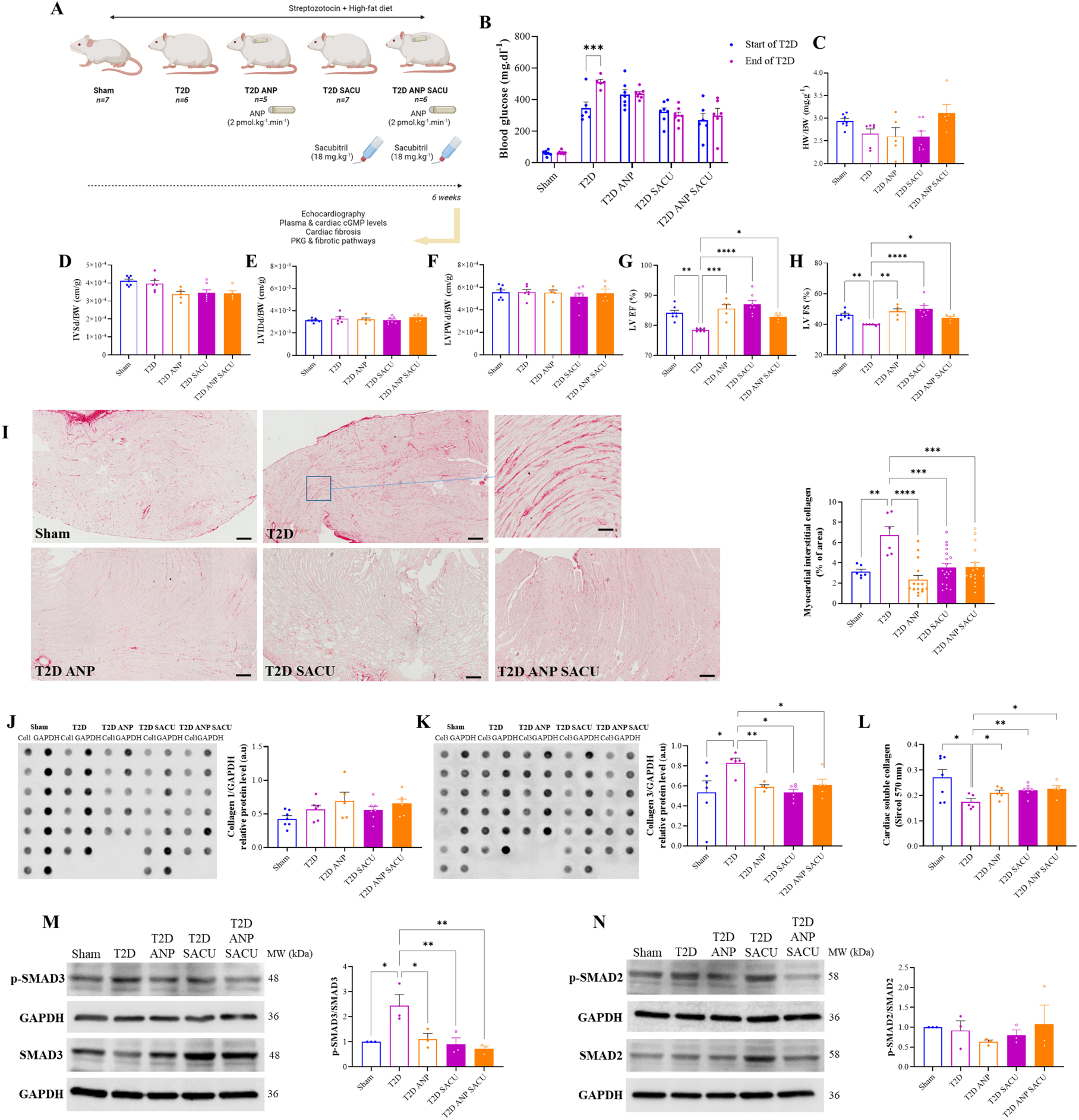
Elevating natriuretic peptides alleviates T2D-induced myocardial fibrosis through the modulation of collagen turnover and SMADs signaling. **A,** Streptozotocin injections and a high-fat diet were used to generate Type 2 Diabetes (T2D) in male rats. ANP (2 pmol.Kg^-1^.min^-1^) was delivered subcutaneously via osmotic pumps, and sacubitril (18 mg.Kg^-1^.day^-1^) was provided orally. At six weeks, echocardiography examined cardiac function, followed by euthanasia for histological and molecular investigations of hearts and plasma. To account for differences in rat size, echocardiographic data were adjusted by dividing by body weight. **B,** Blood glucose in Sham, T2D, T2D ANP, and T2D MANP. **C,** Mean heart weight over body weight ratio (HW/BW), **D,** mean interventricular septum thickness at end-diastole, **E,** mean left ventricular internal dimension at end-diastole, **F,** mean left ventricular posterior wall thickness at end-diastole, **G,** mean left ventricular ejection fraction, and **H,** mean left ventricular fractional shortening in all groups. **I,** Representative heart sections for all groups showing Sirius red staining with respective quantifications of analyzed section areas. **J, K,** Antibody arrays for cardiac collagen I and III using GAPDH as internal control with respective quantifications for all groups. **L,** Cardiac soluble collagen for all groups using Sircol assay. **M, N,** Representative western blots and densitometric analyses of phosphorylated and total SMAD2/3 (each with GAPDH) for all groups. T2D: type 2 diabetes; ANP: atrial natriuretic peptide; MANP: M-atrial natriuretic peptide. All quantitative data are reported as mean and individual data points ± SEM. Normal distribution of the values was checked by the Shapiro-Wilk test and equal variance was checked by the Bartlett’s test. Repeated-measures two-Way ANOVA followed by Sidak’s test in (**B**), One-way ANOVA followed by Sidak’s test in (**D, E, F, G, M**) or Holm-Sidak’s test in (**H, J, K, N**), and Kruskal-Wallis followed by Dunn’s test in (**C, I, L**) were performed. \**P*<0.05, \*\**P*<0.01, \*\*\**P*<0.001, and \*\*\*\**P*<0.0001.

### Sacubitril causes cardiac cGMP accumulation in the myocardium of T2D rats

Cardiac and plasma cGMP levels were regarded as critical indicators for assessing peptide bioactivity. Cardiac cGMP levels fell (by 65%) in T2D, but rose back to normal (≈1.5 fold *vs* T2D) with ANP, sacubitril, or the combination. Nonetheless, neither the disease nor the various treatments had any effect on plasma cGMP levels (Figure 5A, 5B). T2D hindered the cardiac cGMP-dependent pathway PKG/VASP, as demonstrated by 60% lower VASP phosphorylation levels. ANP, sacubitril, and the combination, on the other hand, activated this pathway, but the increase was only statistically significant with the latter (*P*=0.17, *P*=0.16, and *P*<0.05, respectively, Figure 5C). These results indicate that sacubitril and ANP similarly activate the cGMP signaling pathways in the myocardium of T2D rats.

**Figure 5.**
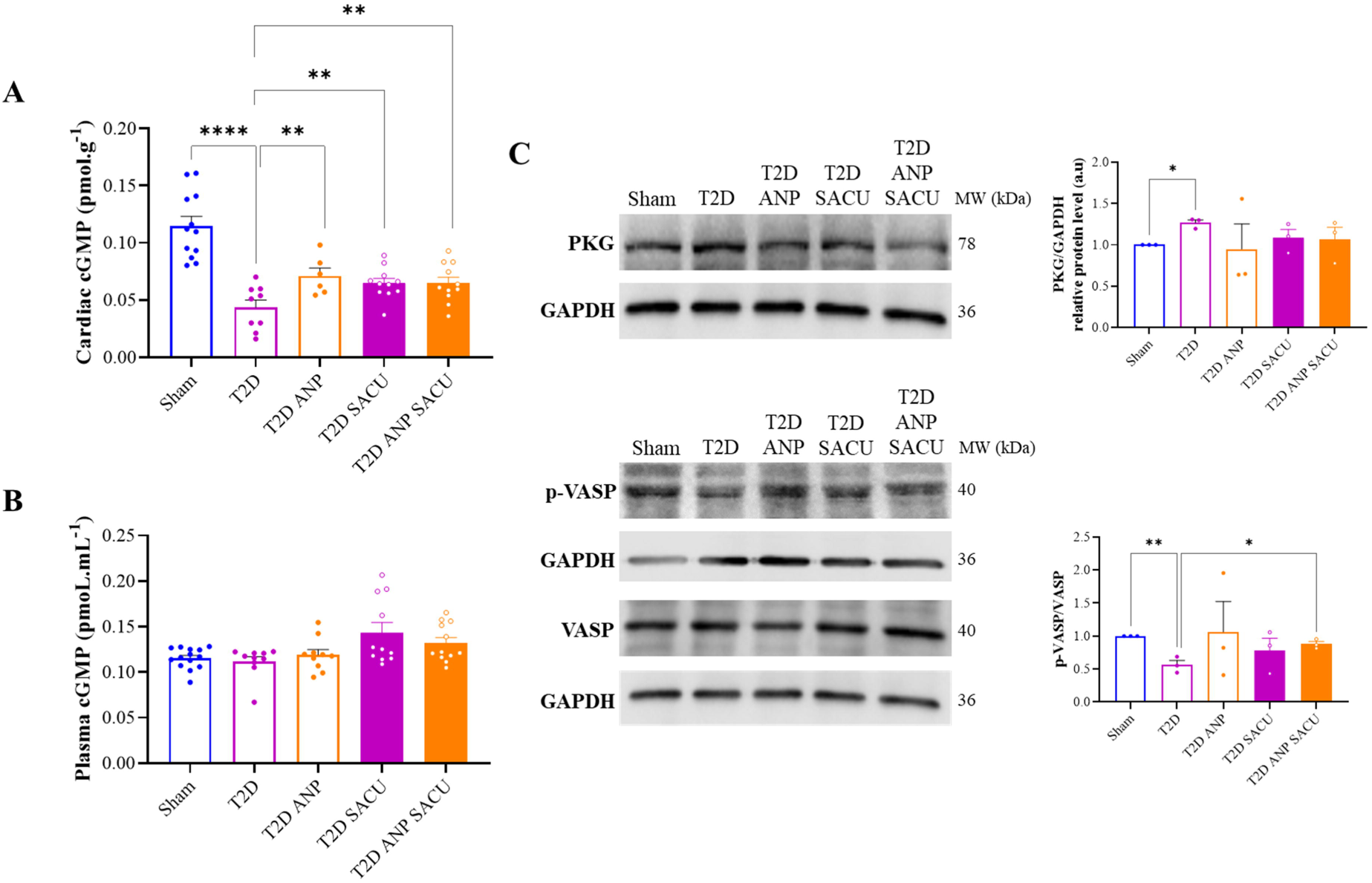
Impact of ANP-enhancing strategies on cardiac cGMP levels. **A, B** Plasma and cardiac cGMP concentrations measured by ELISA in the different rat groups. **C,** Western blots and quantifications of cardiac PKG, total and phosphorylated VASP in the different rat groups, with GAPDH as an internal control (n=3 for each condition). T2D: type 2 diabetes; ANP: atrial natriuretic peptide; MANP: M-atrial natriuretic peptide. All quantitative data are reported as mean and individual data points ± SEM. Normal distribution of the values was checked by the Shapiro-Wilk test and equal variance was checked by the Bartlett’s test. Kruskal-Wallis followed by Dunn’s test in (**B**), One-way ANOVA followed by Sidak’s test in (**A**), and Brown-Forsythe and Welch ANOVA followed by Dunnett’s T3 in (**C**) were performed. \**P*<0.05, \*\**P*<0.01, and \*\*\*\**P*<0.0001.

## Discussion

The main finding of the present study is that myocardial bioavailability of ANP is a major determinant of peptide efficacy in reducing cardiac fibrosis and improving pump function during diabetic cardiomyopathy. We found a balance between peptide elimination and activation of cGMP-dependent signaling pathways that plays a crucial role in the anti-fibrotic effects of ANP on myocardium. Furthermore, our results indicate that reducing myocardial fibrosis is a major therapeutic target of heart failure that can be reached by modulating natriuretic peptides half-life ^30–32^.

Among diabetic experimental models, STZ-induced diabetes reliably replicates diabetes-related features and complications in humans ^33^ notably prominent myocardial fibrosis ^21–26^. Furthermore in the literature, an anti-fibrotic effect of natriuretic peptides has been reported during heart failure ^30–32,34^. Besides, other studies have described a local function of the ANP/GC-A system in moderating the molecular program of cardiac hypertrophy ^35^. In this regard, the enhanced anti-fibrotic effect of the mutated *vs* wild type ANP could account for the former’s superior efficacy in ameliorating cardiac function of STZ-diabetic rats. Cardiac fibrosis is characterized by collagen turnover and phenotype switch leading to increased myocardial stiffness ^36^. Herein, we found an important decrease in collagen type III in the diabetic hearts under MANP along an increase in the soluble collagen. This collagen phenotype fine-tuning might play a crucial role in preserving cardiac performance of the fibrotic heart. In the early stages of heart failure, collagen type III replaces collagen type I leading to cardiomyocyte slippage and fibrotic extracellular matrix. Therefore, the breakdown of collagen type I and the buildup of poorly cross-linked collagen type III are crucial for myocardial fibrosis and ventricular remodeling ^37^. At the molecular level. ANP and TGF-β are known to play counterregulatory roles in cardiovascular physiology and disease. TGF-β acts via its downstream SMAD proteins to mediate fibroblast proliferation, differentiation and collagen deposition, whereas ANP acts as an inhibitor of this pathway ^28,38^. Moreover, collagen assembly is subtly regulated by several structural proteins, in particular Secreted Protein Acidic and Rich in Cysteine (SPARC), that seems to be upregulated in the diabetic heart ^39,40^, stimulated by TGF-β ^41^, and thus might be inhibited by ANP.

Cardiac fibroblasts are the key cellular players in myocardial fibrosis through their high potential to migrate, proliferate and trans-differentiate into myofibroblasts that synthetize ECM components ^42,43^. Several markers have been used to study cardiac fibroblast population in the normal and diseased hearts ^44,45^. Our findings show that both wild-type and MANP similarly inhibit fibroblast activation *in vitro* as evidenced by the reduced collagen production and proliferation. We previously reported that the natriuretic peptides regulate fibroblast activation through the ANP/GC-A/cGMP signaling cascades ^46^. Interestingly, both wild-type and MANP show comparable effects on intracellular cGMP accumulation, cell proliferation, collagen secretion and SMAD activation, indicating that the two peptide isoforms share the same affinity for NPR-A and NPR-C clearance receptors as reported in the literature ^13,14^. Therefore, *in vitro* data cannot explain the difference observed *in vivo* between wild type and mutated peptide on myocardial fibrosis and the upregulation of TGF-β SMAD-dependent pathway ^47^. The explanation is most likely due to reduced clearance of the mutated peptide responsible for enhanced cGMP myocardial concentration together with the upregulation of the cGMP-dependent PKG signaling pathway in MANP vs ANP treated STZ-diabetic rats ^13,14^. The slight increase in plasma cGMP in MANP treated animals might also results from low peripheral peptide degradation. To test the involvement of the coupling between ANP degradation and activation of cGMP-dependent signaling pathway, STZ-diabetic rats were treated with the neprilysin inhibitor sacubitril. Indeed, sacubitril treatment demonstrated similar efficiency to ANP in improving cardiac performance, reducing myocardial fibrosis, and inhibiting the SMAD profibrotic signaling pathway. Furthermore, sacubitril, like ANP, increased myocardial cGMP accumulation and upregulated cGMP-dependent PKG signaling. These findings are consistent with sacubitril’s ability to reduce biomarkers of profibrotic signaling in heart failure ^48^. Of note, sacubitril, contrary to MANP, is not superior to ANP in terms of fibrosis reduction and cardiac function improvement. This might be due to some compensatory clearance pathways such as IDE, other endopeptidases or the NPR-C clearance receptor ^3^. In fact, NPR-C deletion inhibits TGF-β pathway in cardiac fibroblasts leading to reduced myocardial fibrosis ^38^. Nevertheless, the expression and activation pattern of ANP receptors as well as endopeptidase activity remain poorly investigated in diabetic cardiomyopathy and heart disease ^38,49,50^. In view of these multifactorial complexities, more research is required to dissect the pathways underlying the antifibrotic effect of MANP along with ANP. While the cGMP pathway is compromised in patients with heart failure ^4,5,51^, cGMP-enhancing therapies are increasingly showing promising clinical outcomes ^52,53^.

### Limitations

This study presents certain limitations. For a more comprehensive evaluation of MANP’s effects on cardiac fibrosis, longer-term studies should be conducted at advanced stages of heart failure as well as on survival outcomes. Additionally, exploring a range of MANP dosages could provide valuable insights into its safety and effectiveness. While our results demonstrate MANP’s substantial effect on cardiac fibrosis, potential hemodynamic effects of ANP and MANP require further exploration.

## Conclusion

The ANP antifibrotic cardiac effect is related to its myocardial bioavailability which is determined by the rate of peptide degradation. The efficacy of MANP in improving cardiac performance in diabetes appears to be primarily related to its myocardial accumulation and activation of cGMP-dependent PKG signaling pathways. Enhancing ANP tissue accumulation could be an important and promising therapeutic strategy in the management of fibrosis in heart failure.

## Acknowledgment and Funding

This work was jointly funded by the Research Council of Saint-Joseph University of Beirut (Grant #FM348), the CNRS-L Grant Research Program (GRP 2019), and by PHC CEDRE PROJECT N°4782VA.

## Nonstandard Abbreviations and Acronyms

ANP: atrial natriuretic peptide
BacMam: baculovirus transduction of mammalian cells
BNP: brain natriuretic peptide
cGMP: cyclic guanosine monophosphate
EF: ejection fraction
FS: fractional shortening
GAPDH: glyceraldehyde-3-phosphate dehydrogenase
GENIe: genetically encoded nucleotide indicator
IDE: insulin-degrading enzyme
MANP: mutated atrial natriuretic peptide
NP: natriuretic peptides
NPR-C: natriuretic peptide receptor C
PDGFRα: platelet-derived growth factor receptor alpha
PKG: protein kinase G
SACU: sacubitril
SMAD: mothers against decapentaplegic homolog
STZ: streptozotocin
T2D: type-2 diabetes
TGF-β: transforming growth factor beta
VASP: vasodilator-stimulated phosphoprotein
WGA: wheat germ agglutinin

---

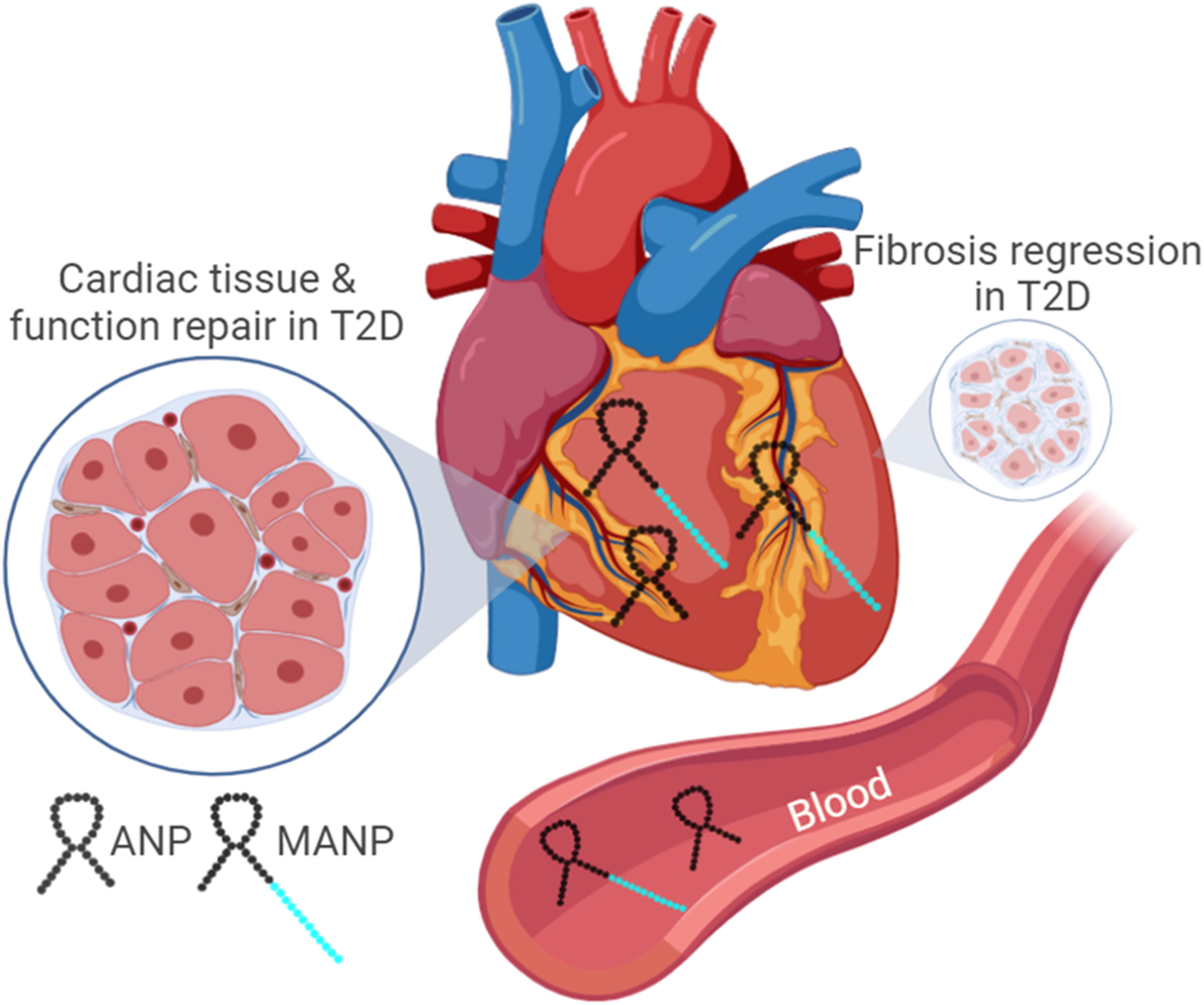

